# Krüppel-like factor 1 acts upstream of the SKN-1/Nrf transcription factors to modulate oxidative stress, lipid homeostasis and longevity

**DOI:** 10.1101/2025.11.22.689963

**Authors:** Jorge Iván Castillo-Quan, Aiden McCarty, Ugne Kurdeikaite, Katherine Gilmore, Arlette Cabral, Justin Mejia, Julia Barrett, Maria La Terza, Emma Johnson, Allison Carroll, T. Keith Blackwell, Jean E. Schaffer, Natalie Moroz

## Abstract

Maintenance of lipid and redox homeostasis are essential for stress resistance and longevity, but the transcriptional networks coordinating these processes remain incompletely understood. In *Caenorhabditis elegans*, the transcription factors SKN-1A/Nrf1 and SKN-1C/Nrf2 mediate distinct stress responses that promote proteostasis, lipid homeostasis, and oxidative stress. Here we identify the Krüppel-like factor KLF-1 as a critical upstream regulator of both SKN-1A and SKN-1C. We show that KLF-1 is required for the oxidative stress resistance and longevity of germline-deficient animals. Genetic interaction studies showed that KLF-1 acts in parallel to the lipogenic regulator Sterol regulatory element-Binding Protein 1 (SBP-1/SREBP1), whereas the related KLF-2 exerts opposing effects on lipid accumulation through SBP-1. Together, these findings place KLF-1 and KLF-2 within a transcriptional network that integrates lipid metabolism, oxidative stress responses, and aging. This work uncovers a conserved regulatory network linking KLFs and SKN-1/Nrf transcription factors in the maintenance of lipid homeostasis and longevity assurance.

## INTRODUCTION

Coordinated gene expression by transcription factors is essential for organismal development, reproduction, and homeostatic maintenance during stress (Himanen et al. 2022; Balasubramaniam et al. 2025; Xie et al. 2025). Exposure to oxidative stress or xenobiotics requires synchronized expression and action of enzymes that neutralize, detoxify and repair cellular damage (Blackwell et al. 2015). Excessive accumulation of certain lipid species can cause stress, lipotoxicity and the onset of disease. However, lipid species vary, and many can produce biologically beneficial effects (Wang and Hu 2017). The continuing rise of obesity and associated metabolic diseases highlights the urgent need to define the transcriptional pathways that govern lipid homeostasis and targets that protect against metabolic dysfunction.

*Caenorhabditis elegans* serve as an invaluable model for elucidating conserved genetic pathways that connect metabolism and homeostatic responses to longevity (Lapierre and Hansen 2012; Ewald et al. 2018). For example, in *C. elegans,* mutations in the *glp-1* Notch gene result in ablation of the germline stem cells (GSC (−)), without affecting the development of the somatic gonad. This results in lifespan extension with increased lipid accumulation (Arantes-Oliveira et al. 2002; O’Rourke et al. 2009). Under physiological conditions, yolk lipoproteins synthesized in intestinal cells are transported to the gonad as a source of nutrients. In the absence of developing oocytes, yolk remains in the pseudocoelomic space or returns to intestinal cells where it stimulates synthesis of triacylglycerides (TAGs) (Lemieux and Ashrafi 2015; Steinbaugh et al. 2015), enhances lipid accumulation (O’Rourke et al. 2009; Steinbaugh et al. 2015), and the activation of several stress response transcription factors, many of which are required for GSC(−) lifespan extension (Kenyon 2010; Antebi 2012; Hansen et al. 2013).

The SKN-1/Nuclear Factor Erythroid 2 (NF-E2)-related factors (Nrfs) family of transcription factors are critical regulators of the oxidative stress response and of lipid metabolism (Hayes and Dinkova-Kostova 2014; Blackwell et al. 2015; Cuadrado et al. 2019). *C. elegans skn-1* gives rise to three functional basic region leucine zipper (bZIP) transcription factors: SKN-1A, SKN-1B, and SKN-1C. SKN-1B is constitutively expressed in a pair of head neurons with no evident mammalian ortholog (Blackwell et al. 2015; Tataridas-Pallas et al. 2021). In contrast, SKN-1A and SKN-1C have clear mammalian orthologs, which mediate defined stress responses (Blackwell et al. 2015; Ruvkun and Lehrbach 2023; Jochim et al. 2025). SKN-1A, the ortholog of Nrf1, is tethered to the endoplasmic reticulum (ER) membrane with most of the protein in the ER lumen (Northrop et al. 2020; Ruvkun and Lehrbach 2023). It is constantly removed from the ER (retrotranslocated) by the ER-associated degradation (ERAD) machinery and targeted for proteasomal degradation. Under conditions of proteasomal stress, such as chemical inhibition or the presence of misfolded cytosolic peptides, SKN-1A/Nrf1 enters the nucleus where it upregulates expression of all proteasome subunit genes. This canonical SKN-1A/Nrf1 stress-response is known as the bounce-back or proteasome recovery response (Northrop et al. 2020; Ruvkun and Lehrbach 2023). Recently, we found that SKN-1A can be activated independently of proteasomal inhibition, at the level of the ER by the monounsaturated fatty acid (MUFA) oleic acid (OA) (Castillo-Quan et al. 2023). OA supplementation results in increased synthesis of triglycerides (TAGs) containing oleic acid and growth of lipid droplets (LDs), both of which enhance ERAD-dependent SKN-1A/Nrf1 activation. As a result, SKN-1A/Nrf1 reshapes the lipid transcriptome, reducing the expression of lipogenic genes while enhancing expression of catabolic genes, in addition to enhancing oxidative stress-related genes, ERAD, and proteasome subunit genes. We termed this response the SKN-1A/Nrf1 lipid homeostatic response to distinguish it from the canonical proteasome recovery response and to highlight its primary role in limiting excessive lipid accumulation (Castillo-Quan et al. 2023).

In contrast, SKN-1C, orthologous to Nrf2, is cytoplasmic and responds to reactive oxygen species (ROS) and xenobiotics (Blackwell et al. 2015). The mitogen-activated kinase (MAPK) signaling pathway phosphorylates and activates SKN-1C, which translocates to the nucleus, and promotes the expression of phase II detoxification genes such as glutathione s-transferase (*gst-4*) (Inoue et al. 2005; Blackwell et al. 2015), which serves as a transcriptional reporter for both SKN-1A/Nrf1 and SKN-1C/Nrf2 (Castillo-Quan et al. 2023).

Interestingly, the Krüppel-like factor (KLF) family of transcription factors are critical regulators of lipid metabolism and phase I detoxification. In *C. elegans*, there are three conserved KLF orthologs (*klf-1*, *klf-2*, and *klf-3*) compared to the five KLFs in *Drosophila*, and at least seventeen members in humans (Bieker 2001; McConnell and Yang 2010). These transcription factors possess highly conserved C-terminal C2H2 zinc finger domains, which enable them to function as both activators and repressors by binding CACCC elements and G/C-rich DNA regions, as well as by interacting with intracellular signaling molecules and transcriptional cofactors (Bieker 2001; Brey et al. 2009; McConnell and Yang 2010).

KLFs are expressed in the intestine of *C. elegans*, where they regulate fat metabolism, stress responses, and longevity assurance (Hsieh et al. 2019). KLF-1 and KLF-2 have been linked to lipid metabolism (Hashmi et al. 2008; Ling et al. 2017). However, these studies were performed using mutant strains in which it was recently shown there is compensatory upregulation of other KLFs (Hsieh et al. 2017). KLF-1 translocates to the nucleus in response to ROS and promotes the transcription of phase I xenobiotic detoxification genes (Herholz et al. 2019), which would complement SKN-1C’s role in upregulating phase II detoxification genes (Blackwell et al. 2015). KLF-1 is also regulated by the proteasome (Carrano et al. 2014), whose function is affected by the SKN-1A proteasome recovery response. Despite their functional parallels, the epistatic relationship between SKN-1 and the KLFs, their interaction, has never been explored.

In this study we found that KLF-1 was required for the activation of SKN-1A/Nrf1 and SKN-1C/Nrf2, a function that likely relates to its role in lipid metabolism. KLF-1 was required for lipid accumulation in animals lacking germline, upstream of SKN-1A, and in parallel to the lipogenic master transcription factor SBP-1/SREBP1. These observations indicate that KLF-1, SBP-1 and SKN-1A are part of a transcription factor network acting in concert to modulate lipid homeostasis.

## RESULTS

### KLF-1 is required for SKN-1 activity and longevity of animals lacking a germ line

We previously performed a genome-scale RNAi screen in GSC(−) animals with the SKN-1A and SKN-1C transcriptional reporter *gst-4p*::GFP (Castillo-Quan et al. 2023). Using WormCat for gene category analysis for the lipid-related and proteostatic gene hits (Holdorf et al. 2020), we identified two transcription factors, the master lipogenic Sterol regulatory element-Binding Protein 1 (SBP-1/SREBP1) and KLF-1 (**Fig. 1A, Supplemental Data 1**). We wanted to understand the role of the KLFs on SKN-1. Of the three *klf* genes, knockdown of *klf-1* and *klf-2* from the L1 larval stage to day one of adulthood did not show any obvious defective developmental phenotypes (**Fig. S1A-C**). Knockdown of *klf-3* led to several developmental defects (**Fig. S1A-B**), including a smaller size at day one of adulthood (**Fig. S1C**), impeding further analysis.

**Figure 1:**
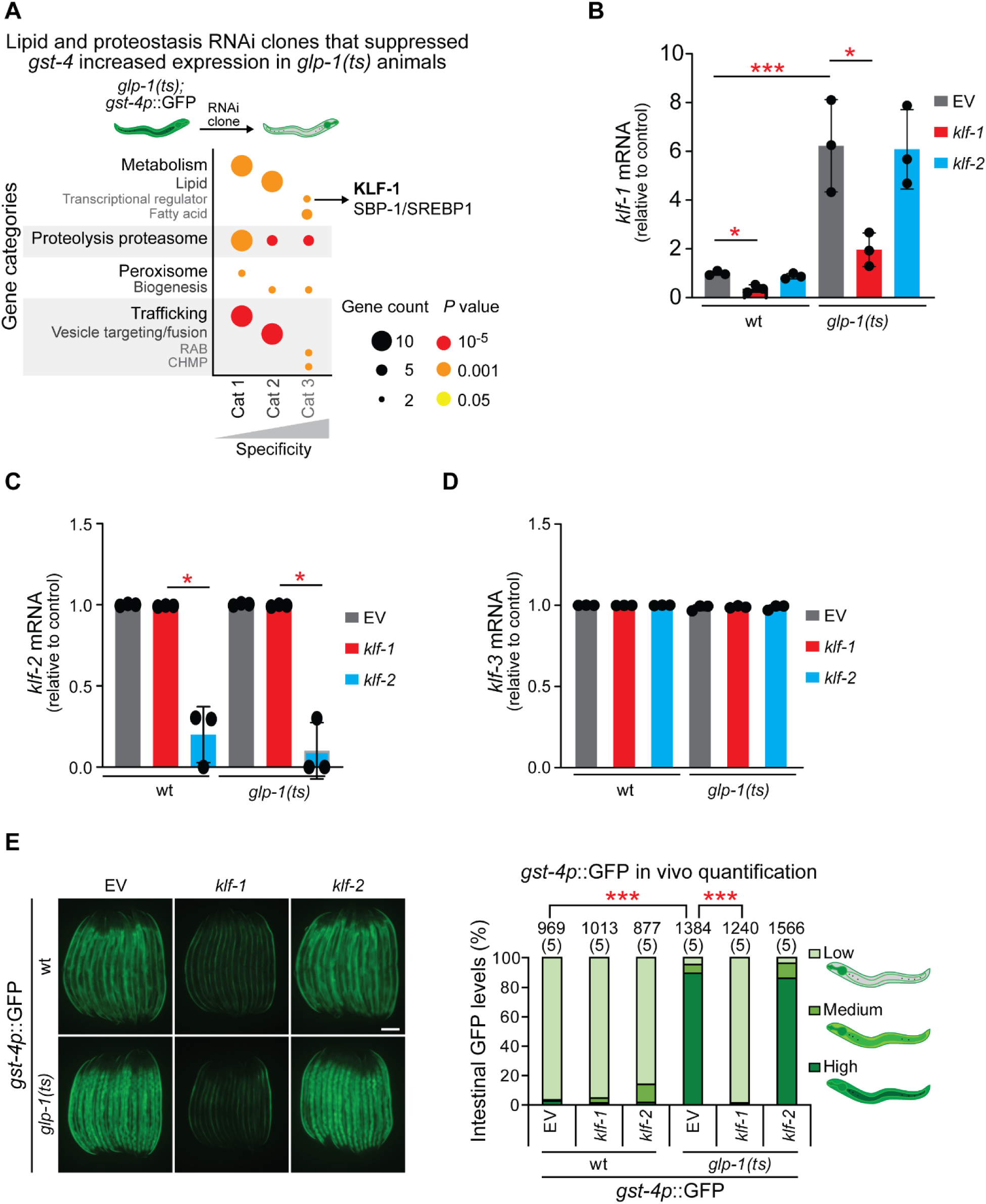
*klf-1* is a positive regulator of SKN-1 transcription. **(A)** Gene categories analyzed by WormCat (Holdorf et al. 2020) of previously identified lipid and proteostasis RNAi clones that suppressed expression of the SKN-1A and SKN-1C transcriptional reporter *gst-4* in germline stem cell ablated animals (GSC(−) or *glp-1(ts)*) (Castillo-Quan et al. 2023). Two transcription factors involved in lipid metabolism were identified including KLF-1, one of three KLF transcription factors. *P* values obtained with Fisher’s exact test with Bonferroni correction for multiple comparisons. See data file S1 for details on genes included in analyses. **(B)** GSC(−) animals have higher expression of endogenous *klf-1* mRNA levels than wild type animals, which are effectively and specifically knocked down by *klf-1* RNAi. ANOVA with Tukey multiple comparisons test. Error bars represent standard deviation. **(C)** *klf-2* mRNA levels are unaffected by GSC(−), but significantly and specifically lowered by *klf-2* RNAi. ANOVA with Tukey multiple comparisons test. Error bars represent standard deviation. **(D)** *klf-3* mRNA is unchanged by GSC(−) and unaffected by knockdown of *klf-1* or *klf-2*. **(E)** *klf-1* knockdown decreases *gst-4p::*GFP expression. Representative images (left) and quantification (right; see data S2 for details). Scale bar, 200 μm. Numbers above bars denote sample size (biological replicates). Pairwise Chi-square tests with False Discovery Rate (FDR) correction (via Benjamini-Hochberg) for multiple comparisons. \**P* < 0.05, \*\**P* < 0.01, ***p < 0.0001.

Both KLF-1 and KLF-2 are expressed during larval and adult stages, with higher expression during larval stages (Hashmi et al. 2008; Zhang et al. 2009; Ling et al. 2017). We explored whether *klf-1* mRNA levels differed in GSC(−) animals and the efficiency of RNAi against the *klf*s. Animals lacking GSC(−) showed increased expression of *klf-1* mRNA, with a significant reduction of expression after *klf-1* RNAi, in both wildtype and GSC(−) animals (**Fig. 1B**). Importantly, RNAi against *klf-1* did not change *klf-2* or *klf-3* mRNA levels. Likewise, RNAi against *klf-2* did not cross impact or change the expression of the remaining two *klf* genes (**Fig. 1C-D**). Thus the RNAi against the KLFs are gene specific and GSC(−) animals show increased expression of KLF-1.

Next, we directly compared the effect of knocking down *klf-1* and *klf-2* on the increased *gst-4* expression in GSC(−) animals. We found that RNAi against *klf-1*, but not *klf-2*, significantly reduced *gst-4p*::GFP reporter expression and *gst-4* mRNA levels (**Fig. 1E**, **Supplemental Data 2**, and **Fig. S1C**). Thus, KLF-1 is required for the expression of the SKN-1 transcriptional target *gst-4* in GSC(−) animals.

SKN-1 activity is required for the longevity of GSC(−) animals (Steinbaugh et al. 2015; Wei and Kenyon 2016; Castillo-Quan et al. 2023). We therefore explored whether knocking down *klf-1* or *klf-2* would impact lifespan. While neither changed the lifespan of wild type animals, the longevity of GSC(−) animals was completely abolished by *klf-1* knockdown, and partially by *klf-2* (**Fig. 2A** and **Table S1**). Interestingly, knockdown of *sbp-1*, the second transcription factor that came out of our screen (**Fig. 1A**), which reduced *gst-4* expression in GSC(−) animals similarly to *klf-1* (**Fig S2A**), only partially suppressed the GSC(−)-mediated lifespan extension (**Fig. 2B** and **Table S1**). Thus, KLF-1 is an essential mediator of the longevity effects of germline loss.

**Figure 2.**
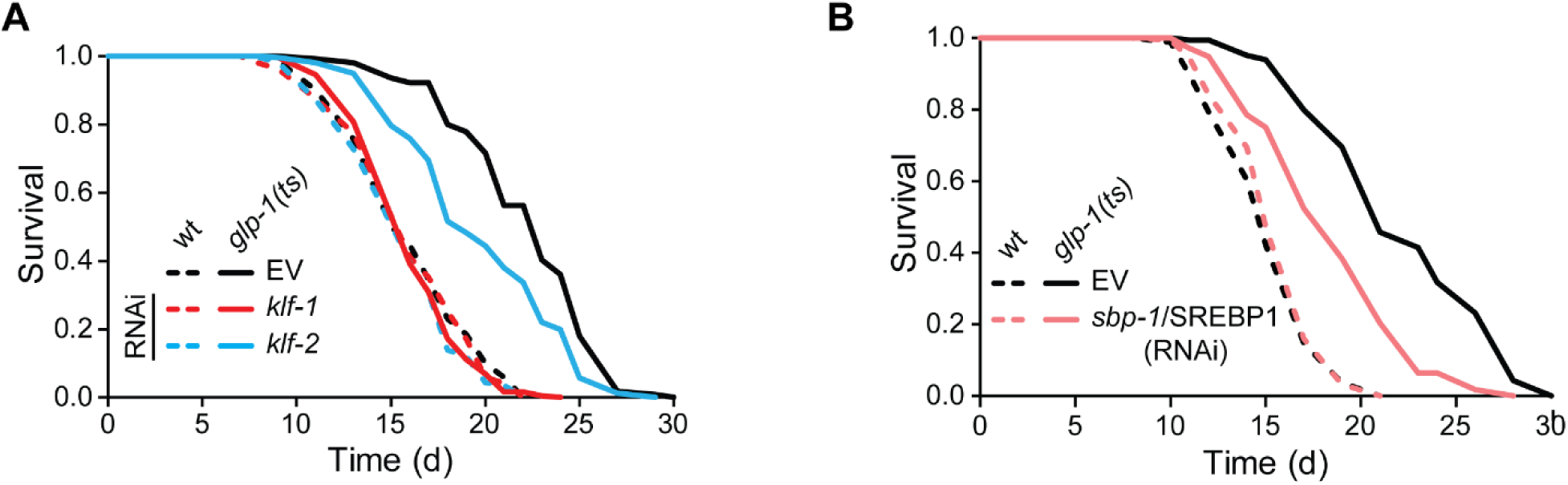
KLF-1 is essential for germline loss-induced longevity. **(A)** *klf-1* knockdown eliminates GSC(−) mediated lifespan extension, while *klf-2* knockdown reduces but does not eliminate longevity (see Table S1 for replicates and statistics). **(B)** Knockdown of *sbp-1*/SREBP1 partially reduces the lifespan extension of GSC(−) animals (see Table S1 for replicates and statistics).

### KLF-1 regulates SKN-1A and SKN-1C activation and response to oxidative stress

To differentiate the role of the KLFs on SKN-1A and SKN-1C, we looked at intestinal nuclear localization of isoform specific protein reporters, which are both nuclear localized in GSC(−) animals (Steinbaugh et al. 2015; Castillo-Quan et al. 2023). We found that *klf-1* knockdown reduced nuclear localization of both SKN-1A/Nrf1 and SKN-1C/Nrf2 in GSC(−) animals, comparable to control levels (**Fig 3A-B**, **Supplemental Data 2**). We analyzed whether KLF-1 mediated this effect by altering the mRNA expression of *skn-1* but did not detect a correlative effect (**Fig. S3A-B**). Instead we detected a specific upregulation of *skn-1* and *skn-1a* mRNA levels in GSC(−) animals subjected to *klf-1* RNAi, a potentially compensatory effect. Thus, KLF-1 likely activates SKN-1A/Nrf1 and SKN-1C/Nrf2 by regulating upstream processes.

**Figure 3.**
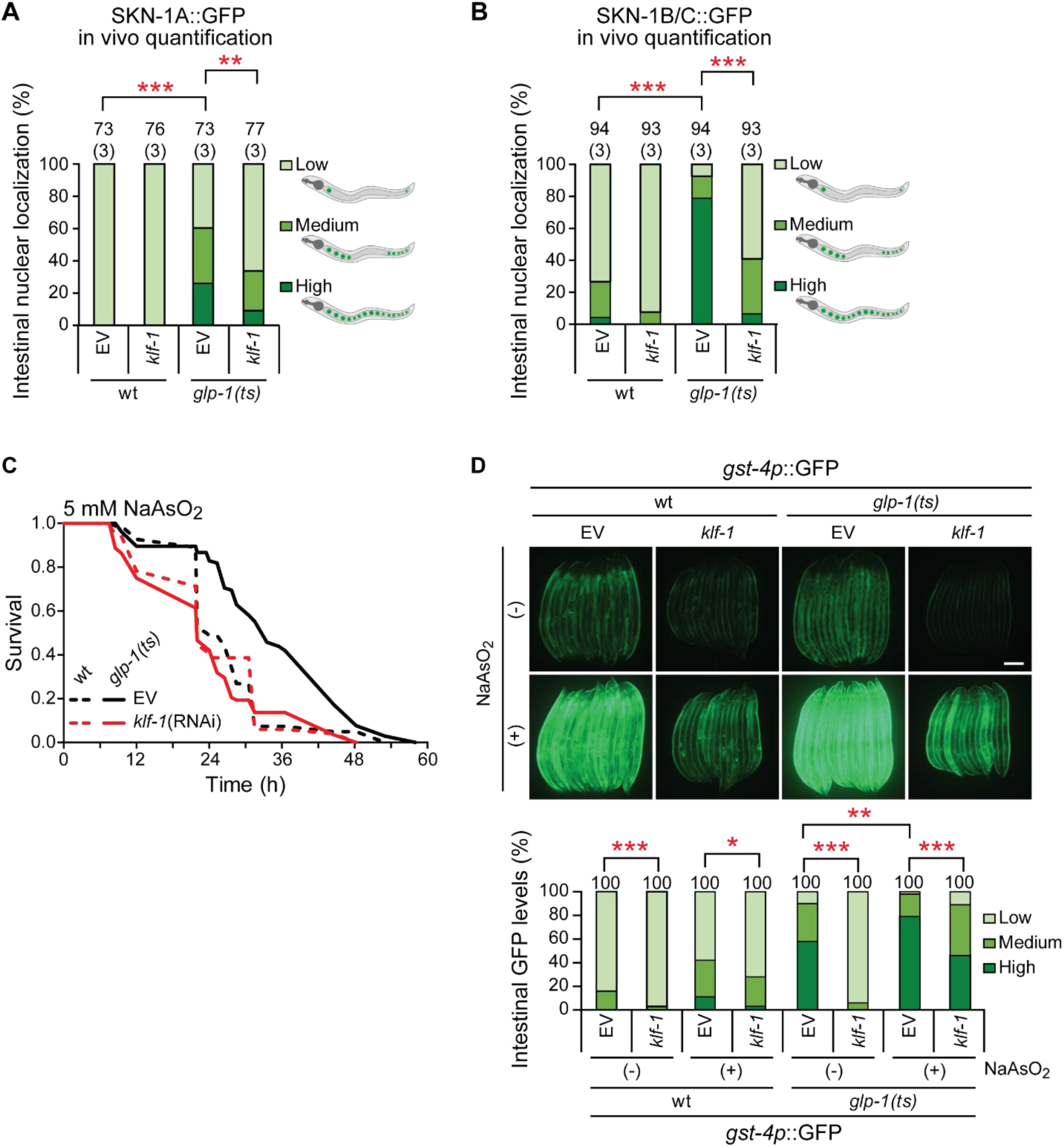
KLF-1 is required for SKN-1A and SKN-1C nuclear localization and oxidative stress resistance. **(A-B)** SKN-1A and SKN-1B/C nuclear localization are reduced by *klf-1* knockdown. Animals were classified (as depicted) and quantified (bar chart). See data file S2 for details. **(C)** Knockdown of *klf-1* abrogates oxidative stress resistance mediated by *glp-1* (see Table S1 for replicates and statistics). **(D)** *klf-1* knockdown eliminates *gst-4* expression induced by sodium arsenite (NaAsO2). Representative fluorescent images (top) and quantification (bottom). Scale bar, 200 μm. **(A/B/D)** Numbers above bars denote sample size (biological replicates). See data file S2 for details. Pairwise Chi-square test with FDR correction for multiple comparisons. \**P* < 0.05, \*\**P* < 0.01, ***p < 0.0001,

To better understand how KLF-1 activates SKN-1A/Nrf1 and SKN-1C/Nrf2, we looked at the role of the KLFs in well-established SKN-1A and SKN-1C mediated activities: the oxidative stress response, the proteasomal recovery response and the lipid homeostasis response. We first exposed animals to the oxidative stress inducer sodium arsenite (NaAsO_2_) and evaluated their survival. As previously reported (Steinbaugh et al. 2015; Castillo-Quan et al. 2023), GSC(−) animals showed increased resistance to oxidative stress in comparison to wild type animals. Interestingly, while *klf-1* knockdown did not increase sensitivity to oxidative stress in wild type animals, it completely abolished the oxidative stress resistance of GSC(−) animals (**Fig 3C** and **Table S1**). We evaluated the SKN-1 transcriptional response to NaAsO_2_-induced oxidative stress by analyzing the GFP levels of the *gst-4p*::GFP reporter. *klf-1* knockdown dampened the NaAsO_2_-mediated increased *gst-4* reporter expression independently of genetic background (**Fig 3D**). Thus, KLF-1 modulates oxidative stress resistance by activating the SKN-1C/Nrf2 transcriptional response.

### KLF-1 modulates the SKN-1A lipid homeostatic response, but not the proteasome recovery response

SKN-1A mediates two homeostatic stress responses: the proteasome recovery response and the lipid homeostatic response (**Fig. 4A**) (Northrop et al. 2020; Castillo-Quan et al. 2023; Ruvkun and Lehrbach 2023). Under steady state conditions, SKN-1A is retrotranslocated out of the ER, ubiquitinated and targeted to the proteasome for degradation (Northrop et al. 2020; Ruvkun and Lehrbach 2023). Alterations that inhibit proteasomal activity, such as cytoplasmic protein aggregates or chemical inhibition with bortezomib, stabilize retrotranslocated SKN-1A, and increase the expression of SKN-1A target genes including all of the proteasomal subunit genes. This is known as the proteasome recovery response (Northrop et al. 2020; Ruvkun and Lehrbach 2023).

**Figure 4.**
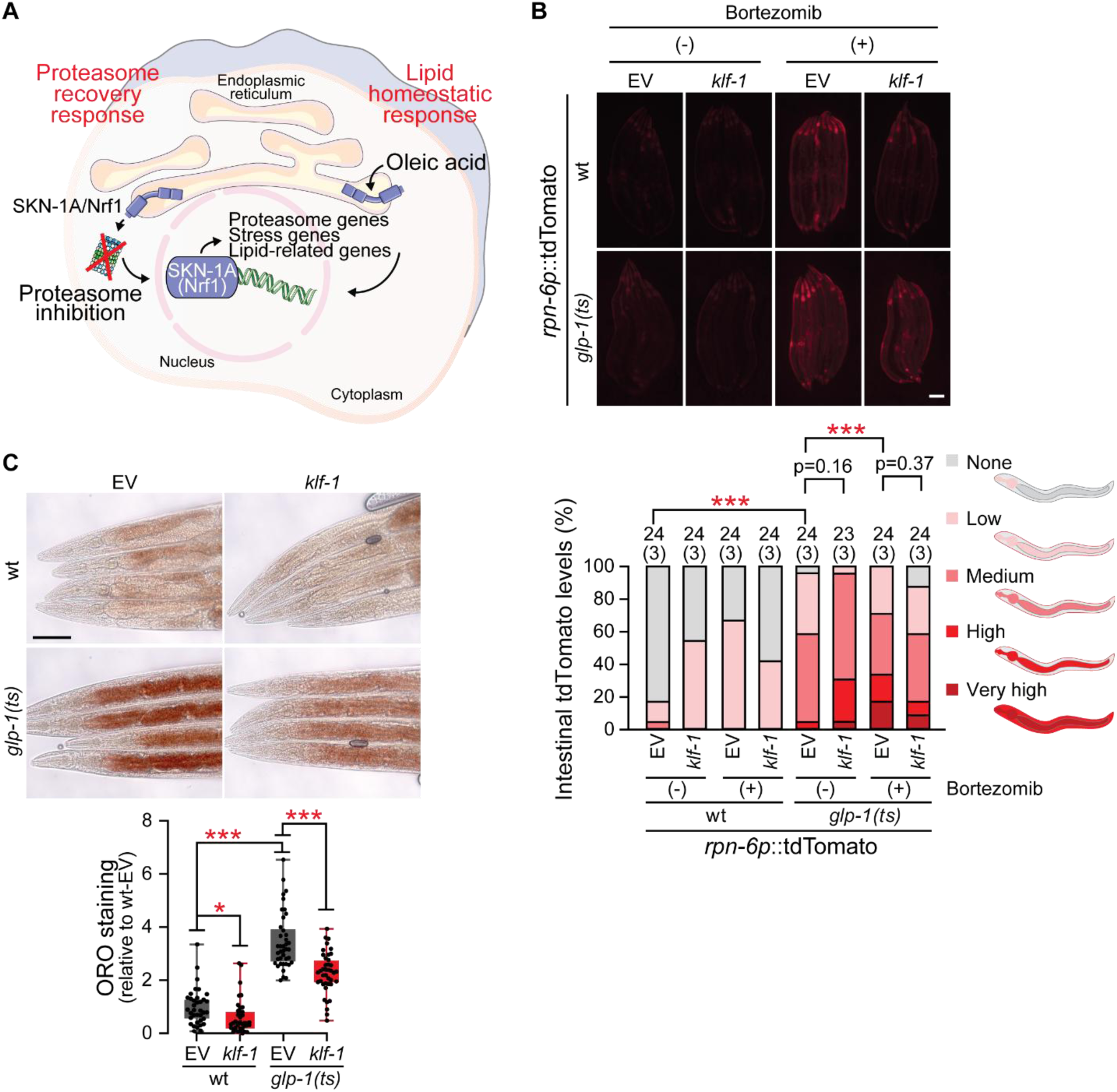
KLF-1 modulates the SKN-1A lipid homeostatic response. **(A)** Diagram showing the two homeostatic stress-response mechanisms in which SKN-1A is transcriptionally activated: the proteasome recovery response and the lipid homeostasis response **(B)** KLF-1 has no modulatory effect on the proteasome recover response. *rpn-6* proteasome subunit reporter increases expression after proteasome inhibition by bortezomib but does not significantly change upon *klf-1* knockdown. Representative images (top) and quantification (bottom). Scale bar, 200 μm. Numbers above bars denote sample size (biological replicates; see data file S2 for details). Pairwise Chi-square test with FDR correction for multiple comparisons. **(C)** KLF-1 is required for the lipid accumulation phenotype in GSC(−) animals as measured by fixed oil red-O (ORO) staining. Representative images (top) and quantification (bottom). Scale bar, 200 μm. \**P* < 0.05, \*\**P* < 0.01, and \*\*\**P* < 0.001.

We first assessed the endogenous mRNA levels of *rpn-6*, a proteasomal subunit, which is known to be increased in GSC(−) animals (Vilchez et al. 2012; Steinbaugh et al. 2015), in a SKN-1A-dependent manner (Castillo-Quan et al. 2023). The increased expression of *rpn-6* mRNA, in GSC(−) animals, was unaffected by *klf-1* knockdown (**Fig. S4A**). We next assessed the role of KLF-1 in the proteasome recovery response by analyzing the fluorescence of the proteasome reporter *rpn-6p*:tdTomato, which is enhanced in GSC(−) animals (Vilchez et al. 2012; Castillo-Quan et al. 2023). Proteasomal inhibition by bortezomib increased *rpn-6* expression independently of the genetic background, while *klf-1* knockdown did not have a significant effect (**Fig. 4B**). We assessed the proteasome recovery response using a second proteasome reporter *rpn-12p*::GFP (Castillo-Quan et al. 2023), and again did not detect a significant effect on the increased proteasome reporter expression after bortezomib exposure, upon *klf-1* knockdown (**Fig. S4B**). The increased expression of *rpn-12* mRNA levels in GSC(−) animals was also unchanged by *klf-1* knockdown (**Fig. S4C**). Thus, KLF-1 has no effect on the increased expression of proteasome genes in GSC(−) animals, or the SKN-1A/Nrf1 proteasome recovery response.

The second response mediated by SKN-1A/Nrf1 is the lipid homeostasis response (**Fig. 4A**). Increased synthesis or accumulation of OA at the ER membrane activates SKN-1A, which increases the expression of lipid catabolism genes, while decreasing expression of lipogenic genes (Castillo-Quan et al. 2023). GSC(−) animals have increased OA-dependent intestinal steatosis and thereby increased SKN-1A activity (O’Rourke et al. 2009; Castillo-Quan et al. 2023). We measured triglyceride levels, the only neutral lipid in *C. elegans,* by fixed oil red O (ORO) staining (Watts and Ristow 2017; Castillo-Quan et al. 2023). In agreement with previous reports, GSC(−) animals showed marked increased levels of ORO staining in comparison to wild type animals. Interestingly, knockdown of *klf-1* significantly reduced ORO staining in both wild type and GSC(−) animals (**Fig. 4C**). Thus, KLF-1 likely acts on SKN-1A/Nrf1 by modulating the lipid accumulation that activates the SKN-1A/Nrf1 lipid homeostatic response.

### KLF-1 modulates lipid metabolism upstream of SKN-1A/Nrf1 and in parallel to SBP-1/SREBP1

To understand how KLF-1 and SKN-1A interact to regulate lipid homeostasis, we looked at lipid accumulation in animals specifically lacking *skn-1a* (Lehrbach and Ruvkun 2016; Castillo-Quan et al. 2023). Loss of SKN-1A results in loss of lipid catabolism and increased lipid synthesis, resulting in intestinal steatosis (Castillo-Quan et al. 2023). We found that knocking down *klf-1* reduced lipid accumulation in both wild type and *skn-1a* null animals (**Fig. 5A**). Thus, the effects of KLF-1 on lipid metabolism are likely independent of SKN-1A.

**Figure 5.**
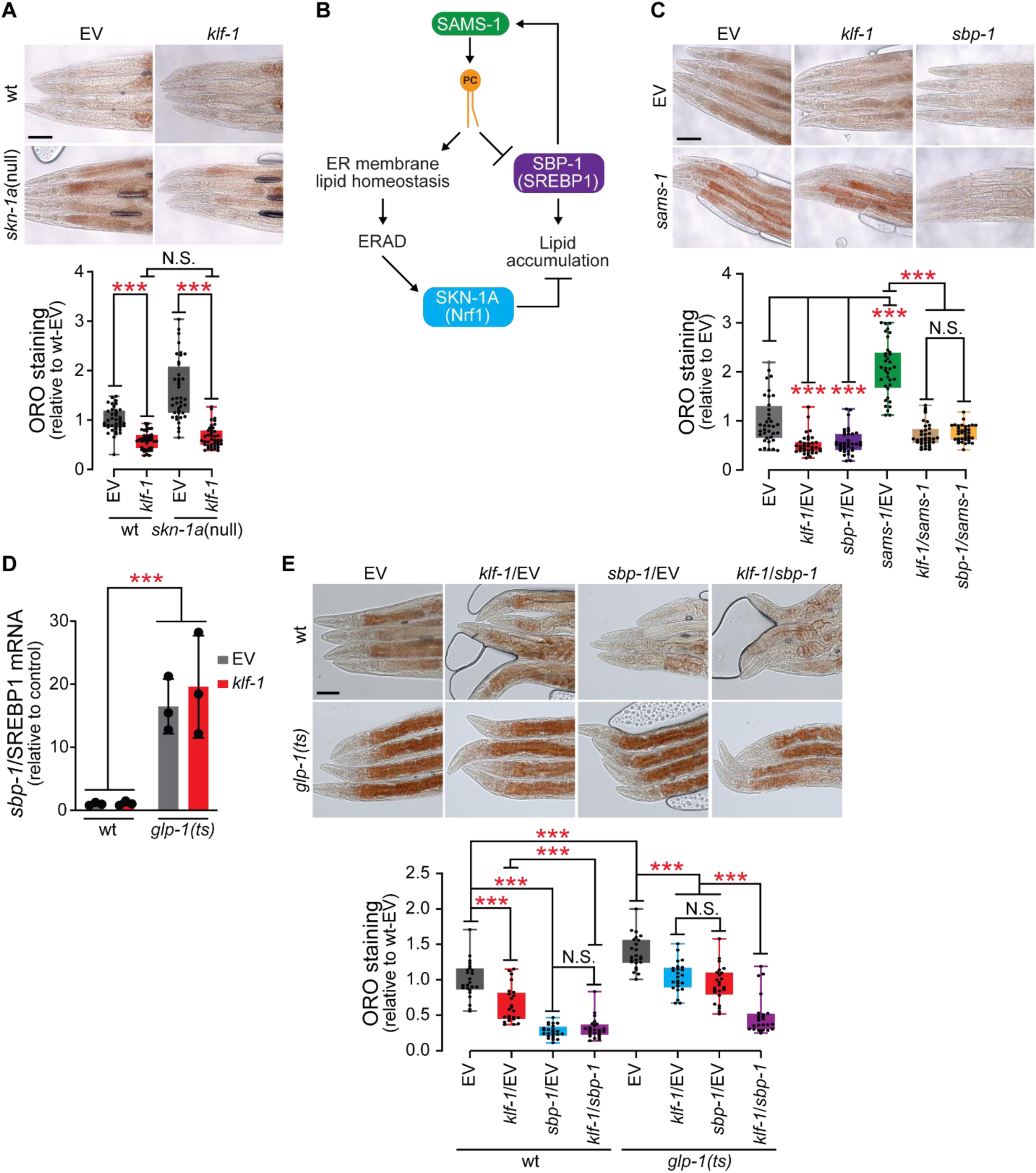
KLF-1 modulates lipid accumulation upstream of SKN-1A and in parallel to SBP-1. **(A)** Knockdown of *klf-1* reduces triglyceride accumulation (ORO staining) in *skn-1a* null animals. Representative images (top) and quantification (bottom). ANOVA with Tukey multiple comparisons test. Scale bar, 200 μm. **(B)** Diagram showing the effect of phosphatidylcholine (PC) synthesis inhibition. On the one hand it impairs SKN-1A/Nrf1 activation by interfering with ERAD efficiency (Castillo-Quan et al. 2023), and on the other hand it activates SBP-1/SREBP1 in a compensatory feedback mechanism that results in lipogenesis (Walker et al. 2011). **(C)** *klf-1* knockdown reduces intestinal steatosis produced by loss of *sams-1*. Representative images (top) and quantification (bottom). Scale bar, 200 μm. **(D)** *sbp-1* mRNA is significantly increased in GSC(−) animals, but unaffected by *klf-1* knockdown. ANOVA with Tukey multiple comparisons test. (**E**) In GSC(−) animals, simultaneous knockdown of *sbp-1* and *klf-1* have an additive effect on reducing the increased lipid accumulation. Representative images (top) and quantification (bottom). Scale bar, 200 μm. ANOVA with Tukey multiple comparisons test.

We previously found that SKN-1A activation in GSC(−) animals requires phosphatidylcholine (PC) synthesis, as reduced PC impairs ERAD function (Castillo-Quan et al. 2023), which is essential for SKN-1A to be retrotranslocated out of the ER membrane, to move to the nucleus (Ruvkun and Lehrbach 2023). Interestingly, inhibition of PC synthesis activates a compensatory transcriptional response in which SBP-1/SREBP1 is activated, resulting in increased lipid accumulation (Walker et al. 2011). Given that the standard laboratory *C. elegans* diet lacks choline, *C. elegans* rely on the conversion of phosphoethanolamine to phosphocholine in the CDP-DAG pathway through the action of the PMT-1/2 methyltransferases and *S*-adenosyl methionine synthase (SAMS)-1 acting as a methyl donor (Watts and Ristow 2017). Hence, SAMS-1 inhibition blocks SKN-1A activation and activates SBP-1/SREBP1 (**Fig. 5B**) (Walker et al. 2011; Castillo-Quan et al. 2023).

We next explored whether KLF-1 had a role in reduced *sams-1* mediated lipid accumulation. As previously reported, *sams-1* RNAi increased lipid accumulation in an SBP-1/SREBP1-dependent manner (**Fig. 5C**) (Walker et al. 2011; Smulan et al. 2016). Interestingly, *klf-1* knockdown reduced lipid accumulation produced by *sams-1* deficiency, similarly to *sbp-1* (**Fig. 5C**). We explored whether KLF-1 could be transcriptionally regulating SBP-1, reducing *sbp-1* mRNA levels, but we did not find evidence to support this hypothesis (**Fig. 5D** and **Fig S5A**). We next assessed whether SBP-1 and KLF-1 act in the same pathway by simultaneously inhibiting them. In wild type animals, knockdown of *sbp-1* produced a stronger lipid lowering phenotype than *klf-1* knockdown, with simultaneous inhibition unable to further reduce the low lipid phenotype upon *sbp-1* knockdown (**Fig. 5E**). However in GSC(−) animals, which have an enhanced lipid profile, simultaneously knocking down *sbp-1* and *klf-1* produced a stronger lipid lowering effect than inhibiting each individually (**Fig. 5E**). Thus, KLF-1 and SBP-1/SREBP1 likely act in parallel pathways to modulate lipid homeostasis.

### KLF-1 and KLF-2 have opposing effects on lipid accumulation

In both *Drosophila* and mammalian systems, KLF transcription factors can exert antagonistic effects on cellular proliferation and differentiation, with some promoting cell cycle exit while others stimulate continued proliferation, reflecting a conserved principle of functional opposition within the family (McConnell and Yang 2010; Wu et al. 2018). This antagonistic effect extends to metabolic actors and phenotypes (McConnell and Yang 2010; Hsieh et al. 2019). We thus wondered whether KLF-2 influenced lipid accumulation. We knocked down *klf-1* and *klf-2* and observed that while *klf-1* RNAi reduced lipid accumulation, decreasing *klf-2* increased lipid accumulation, as measured by ORO staining (**Fig. 6A**). Interestingly, simultaneous knockdown of *klf-1* and *klf-2* brought the lipid phenotype to levels comparable to WT controls (**Fig. 6A**). Thus, *C. elegans* KLF-1 and KLF-2 exert opposing effects on lipid accumulation.

**Figure 6.**
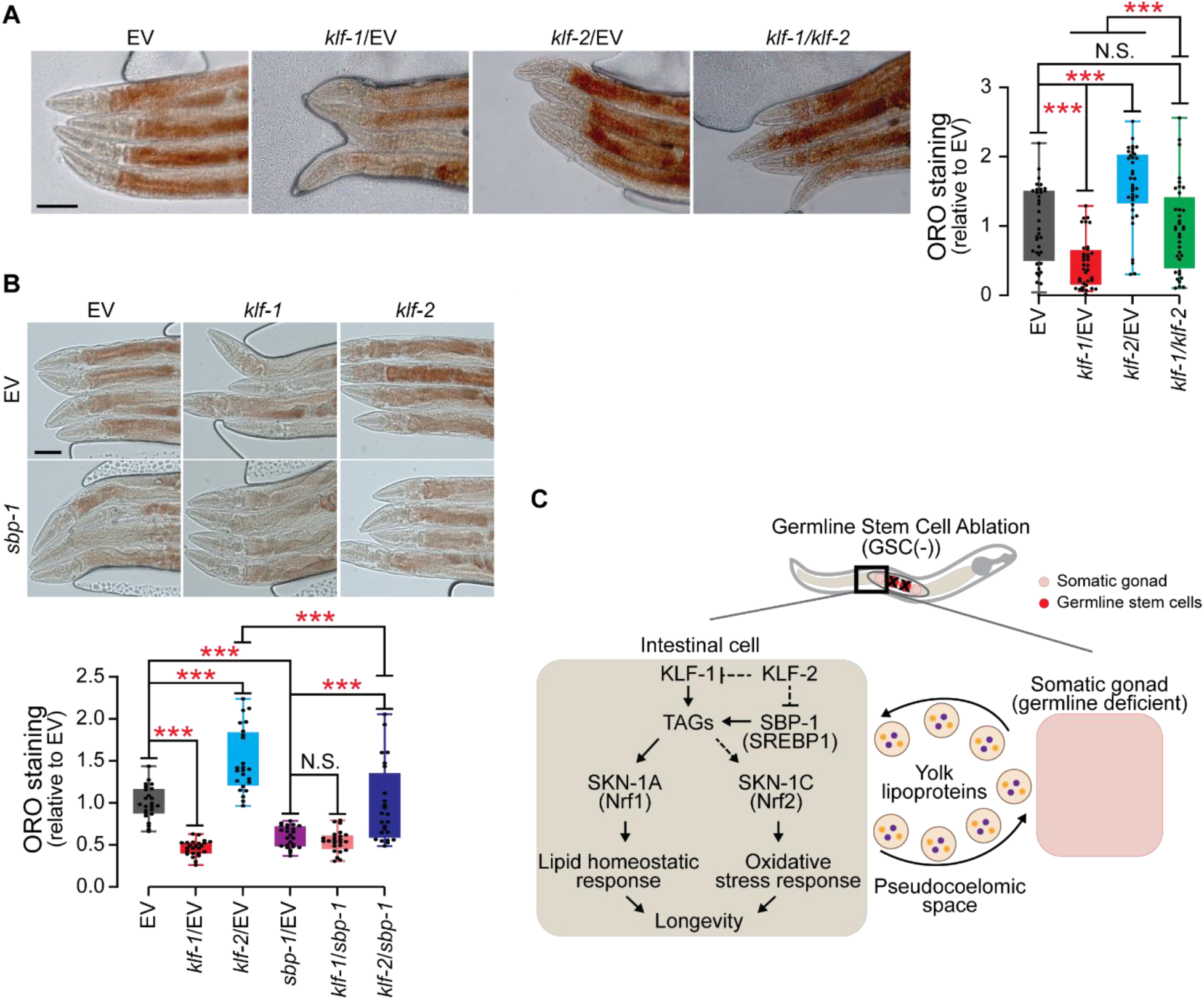
KLF-1 and KLF-2 have opposing roles in lipid metabolism. **(A)** While *klf-1* knockdown reduces lipid accumulation, *klf-2* inhibition increases lipid accumulation. Simultaneous knockdown of *klf-1* and *klf-2* results in triglyceride levels indistinguishable from EV control animals. Representative images (left) and quantification (right). Scale bar, 200 μm. ANOVA with Tukey multiple comparisons test. N.S. = not significant, \*\*\**P* < 0.001. **(B)** *klf-2* knockdown-induced lipid accumulation requires SBP-1/SREBP1. Representative images (top) and quantification (bottom). Scale bar, 200 μm. ANOVA with Tukey multiple comparisons test. N.S. = not significant, \*\*\**P* < 0.001. **(C)** Summary diagram. In GSC(−) animals, the absence of a developing germline results in unused yolk lipoproteins returning to intestinal cells and stimulating lipogenesis. We propose that KLF-1 is an upstream regulator of this lipid accumulation phenotype, driving the activation of SKN-1A/Nrf1 and SKN-C/Nrf2. In contrast, KLF-2 likely acts as a repressor of KLF-1 and SBP-1, and their related processes. KLF-1 therefore integrates lipid metabolism, oxidative stress and longevity in this transcription factor regulatory network.

We showed that KFL-1 likely acts in parallel to the lipogenic SBP-1/SREBP (**Fig. 5E**). Given that KLF-1 and KLF-2 have antagonistic effects on lipid homeostasis, we wondered whether KLF-2 acts through or independently of SBP-1/SREBP1. We confirmed that knockdown of *klf-1* and *klf-2* had a lipid lowering and increasing effect, respectively (**Fig. 6B**). However, simultaneous knockdown of *klf-2* and *sbp-1* abolished the lipid accumulating phenotype of *klf-2* knockdown (**Fig. 6B**). Thus, KLF-2 likely acts as an upstream negative regulator of SBP-1 to modulate lipid metabolism. Together our results show that KLF-1 and KLF-2 are part of a transcriptional regulatory network that maintains lipid homeostasis (**Fig. 6C**). We show that KLF-1 acts in parallel to SBP-1/SREBP1, while KLF-2, which opposes KLF-1s role in lipid metabolism, acts in an inhibitory manner, upstream of SBP-1.

## DISCUSSION

We identified KLF-1 as a central regulator of lipid homeostasis, oxidative stress resistance, and longevity in *C. elegans*. We showed that KLF-1 acts upstream of the conserved transcription factors SKN-1A/Nrf1 and SKN-1C/Nrf2, two functionally distinct isoforms that mediate transcriptional programs modulating proteostatic and lipid metabolic insults, and oxidative stress, respectively. In *C. elegans*, yolk lipoproteins, targeted for the germline, are produced in the intestinal cells and return to these same cells if the animal’s germline is absent and the gonad does not require these nutrients (Lemieux and Ashrafi 2015; Steinbaugh et al. 2015). The returning yolk produces a lipid accumulation phenotype that is dependent on KLF-1 (**Fig. 6C**). We previously showed that this lipid accumulation phenotype is required for SKN-1A/Nrf1 and SKN-1C/Nrf2 activation (Steinbaugh et al. 2015; Castillo-Quan et al. 2023). Hence, in our revised model, KLF-1 acts upstream of these two transcription factors by modulating lipid accumulation (**Fig. 6C**). These findings position KLF-1 as an upstream node linking the lipid metabolic state to activation of stress responses, integrating previously distinct transcriptional modules controlling lipid metabolism, redox, and organismal aging.

### KLF-1 acts on SKN-1C to coordinate the xenobiotic and oxidative stress responses

The Nrf transcription factors SKN-1A and SKN-1C are activated by different cellular cues: SKN-1A/Nrf1 responds to ER lipid changes and proteasomal perturbations (Castillo-Quan et al. 2023; Ruvkun and Lehrbach 2023), while SKN-1C/Nrf2 responds to cytosolic oxidative and xenobiotic stress (An and Blackwell 2003; Blackwell et al. 2015). Our results reveal that both isoforms depend on KLF-1 for their activation in GSC(−) animals, which are characterized by increased lipid accumulation and robust stress resistance, identifying KLF-1 as an integrator of stress responses.

Xenobiotic detoxification involves three sequential phases of detoxification: bioactivation (phase I), conjugation (II), and transport and elimination (III). In *C. elegans*, KLF-1 has been shown to regulate phase I genes, including cytochrome P450 oxidases (CYPs) (Herholz et al. 2019), while SKN-1C/Nrf2 is the master regulator of phase II detoxification genes, including *gst-4* (Inoue et al. 2005; Blackwell et al. 2015). Interestingly, reactive oxygen species (ROS) activate both KLF-1 and SKN-1C/Nrf2 (Hourihan et al. 2016; Hermeling et al. 2022). This would suggest that KLF-1 acts as an upstream modulator of the whole detoxification machinery and oxidative stress response, by directly acting on phase I detoxification genes, and activating SKN-1C/Nrf2 to coordinate phase II detoxification and the redox response. Because KLF-1 depletion did not alter *skn-1* transcript levels, KLF-1 likely affects SKN-1A and SKN-1C through metabolic, redox, or growth signaling pathways known to impact SKN-1A/Nrf1 and SKN-1C/Nrf2 activation (Inoue et al. 2005; Tullet et al. 2008; Paek et al. 2012; Robida-Stubbs et al. 2012; Moroz et al. 2014; Blackwell et al. 2015; Hourihan et al. 2016; Ewald et al. 2018; Meng et al. 2021; Castillo-Quan et al. 2023).

### KLF-1 modulates the SKN-1A lipid homeostatic response

SKN-1A mediates two distinct homeostatic programs: the proteasome recovery response, triggered by proteasomal inhibition, and the lipid homeostatic response, triggered by ER lipid remodeling and accumulation of monounsaturated fatty acids such as oleic acid (Castillo-Quan et al. 2023). Our findings indicate that KLF-1 selectively regulates the lipid branch of this pathway. In GSC(−) animals, lipid redistribution and storage are critical for lifespan extension (Goudeau et al. 2011), and requires both SKN-1A and SKN-1C isoforms (Steinbaugh et al. 2015; Castillo-Quan et al. 2023). Our data suggest that KLF-1 acts as a metabolic rheostat: promoting the synthesis or accumulation of lipid species that trigger the SKN-1A lipid homeostatic response, leading to enhanced lipid catabolism and redox resilience. Thus, KLF-1 provides a mechanistic bridge between lipid metabolism and antioxidant defenses.

The effect of KLF-1 on the SKN-1A lipid homeostatic response suggests that KLF-1 influences the ER lipid environment rather than SKN-1A processing or proteasome activity. Because the ER membrane lipid composition and the function of the ER-associated degradation (ERAD) pathway are critical for SKN-1A activation (Castillo-Quan et al. 2023; Ruvkun and Lehrbach 2023), KLF-1 may regulate lipid synthesis or trafficking processes that tune ER membrane saturation, lipid droplet biogenesis, or phospholipid synthesis, thereby influencing SKN-1A nuclear translocation and transcriptional activity.

### KLF-1 functions in parallel to SBP-1/SREBP-1 to regulate lipid homeostasis

KLF-1’s effects on lipid metabolism are distinct from, though partially overlapping, with those of the lipogenic transcription factor SBP-1/SREBP-1. While *sbp-1* knockdown reduced lipid levels and partially suppressed GSC(−)-mediated lifespan extension, *klf-1* knockdown completely abolished its longevity. Epistasis analysis showed that simultaneous inhibition of *klf-1* and *sbp-1* produced additive effects in GSC(−) animals, consistent with parallel or intersecting pathways. Given that *klf-1* depletion did not alter *sbp-1* transcript levels, these two transcription factors likely act independently to modulate lipid metabolic flux. SBP-1 primarily drives fatty acid and TAG synthesis (Watts and Ristow 2017), whereas KLF-1 may regulate complementary processes such as fatty acid desaturation, lipid droplet turnover, or phospholipid remodeling. Thus, KLF-1 and SBP-1 form a dual regulatory module that balances anabolic and catabolic lipid processes to maintain homeostasis.

### Opposing roles of KLF-1 and KLF-2 in lipid metabolism

We found that *klf-1* knockdown reduced lipid accumulation, while *klf-2* knockdown increased it, and combined inhibition restored neutral lipid levels to baseline. This functional antagonism suggests that KLF-1 and KLF-2 exert opposing regulatory effects on shared lipid metabolic targets or processes. KLF-1 and KLF-2 have each been identified as negative regulators of lipid accumulation (Hashmi et al. 2008; Brey et al. 2009; Zhang et al. 2009; Zhang et al. 2013; Ling et al. 2017; Wang et al. 2023). However, the protocols used significantly differ from ours. We knocked down gene expression starting at the L1 stage and assessed lipid profiles 72 hours later at the day-one adult stage. Previous protocols knocked down *klf-1* at the young adult or L4 stage by soaking animals in dsRNA for 24hr under fasting conditions, followed by Sudan black staining, Nile Red staining, or refeeding for 24 hours followed by ORO staining (Hashmi et al. 2008; Brey et al. 2009; Wang et al. 2023). It remains a possibility that short-term knockdown yields different responses, or that KLFs have developmental stage specific effects. We consistently observed that KLF-1 is a positive regulator of lipid metabolism and KLF-2 a negative regulator.

### A conserved regulatory network linking KLFs and Nrf transcription factors

Antagonism between KLF paralogs is a conserved feature in metazoans: in mammalian systems, distinct KLFs either promote lipid storage or lipid oxidation depending on tissue context (McConnell and Yang 2010; Hsieh et al. 2019). Meanwhile, the integration of KLF and Nrf transcription factors defines regulatory architecture that connects metabolic and stress-responsive pathways. In mammals, Nrf1 and Nrf2 govern proteostasis and antioxidant gene expression, while several KLFs, including KLF2, KLF4, and KLF15, modulate lipid and redox metabolism (Hsieh et al. 2019). Our data suggest that the cooperation between these transcription factor families is evolutionarily conserved. Specifically, KLF-1 in *C. elegans* may represent a functional analog of mammalian KLF15, which regulates lipid utilization and redox homeostasis in the liver and muscle.

### Concluding remarks

In summary, our findings establish KLF-1 as a key metabolic regulator that links lipid metabolism, oxidative stress resistance, and longevity in *C. elegans*. Acting upstream of SKN-1A/Nrf1 and SKN-1C/Nrf2, KLF-1 modulates lipid accumulation and detoxification gene expression, while functioning in parallel to SBP-1/SREBP-1. Together with the antagonistic activity of KLF-2, these interactions define a transcriptional network by which lipid metabolic stress sensed by the KLFs primes the activation of Nrf transcriptional programs to restore homeostasis. The conservation of KLF and Nrf signaling across metazoans suggests that similar mechanisms may coordinate metabolic and redox balance in higher organisms, with potential implications for understanding metabolic dysfunction, aging, and related diseases.

## Methods

### *C. elegans* culture and maintenance

*C. elegans* strains were maintained at 15°C or 20°C on nematode growth medium (NGM) plates seeded with *Escherichia coli* (OP50) using standard techniques (Brenner 1974). To produce synchronized populations of worms, day-one adults were bleach-synchronized using M9 buffer to wash worms of the plate and adding bleach and 5M NaOH. After obtaining unhatched embryos through continuous vertexing, the sample was washed at least three times with M9 buffer (centrifugation at 700 rpm for 1 min). The remaining *C. elegans* embryos were resuspended in 10 mL of M9 buffer and left to hatch and arrest at the L1 stage for 24 hours at 15°C or 20°C. L1 animals were seeded onto the appropriate bacteria and placed again at the appropriate experimental temperature. The *skn-1a*(null) strain experiences a slight developmental delay therefore *skn-1a*(null) L1s were seeded 6 hours before N2 L1s. Strains containing *glp-1*(bn18) were maintained at 15°C (permissive temperature), and the following temperature shift protocol was used (Steinbaugh et al. 2015; Castillo-Quan et al. 2023). Day-1 adult worms grown at 15°C were bleach-synchronized. The *C. elegans* embryos were left to hatch and arrest at the L1 stage for 48 hours at 15°C. L1 animals were seeded onto the appropriate bacteria and placed again at 15°C. Animals were then shifted to the nonpermissive temperature of 25°C (preventing development of the GSC) at the L2 stage and remained at this temperature for the duration of the experiment (Steinbaugh et al. 2015; Castillo-Quan et al. 2023). Given the effect of temperature on many of the phenotypes that we evaluated, non-*glp-1* strains were subjected to the same protocol when compared to *glp-1* strains. In experiments that did not include *glp-1* strains, *C. elegans* were not subjected to this temperature shift protocol. The *C. elegans* strains used in this study are detailed here:

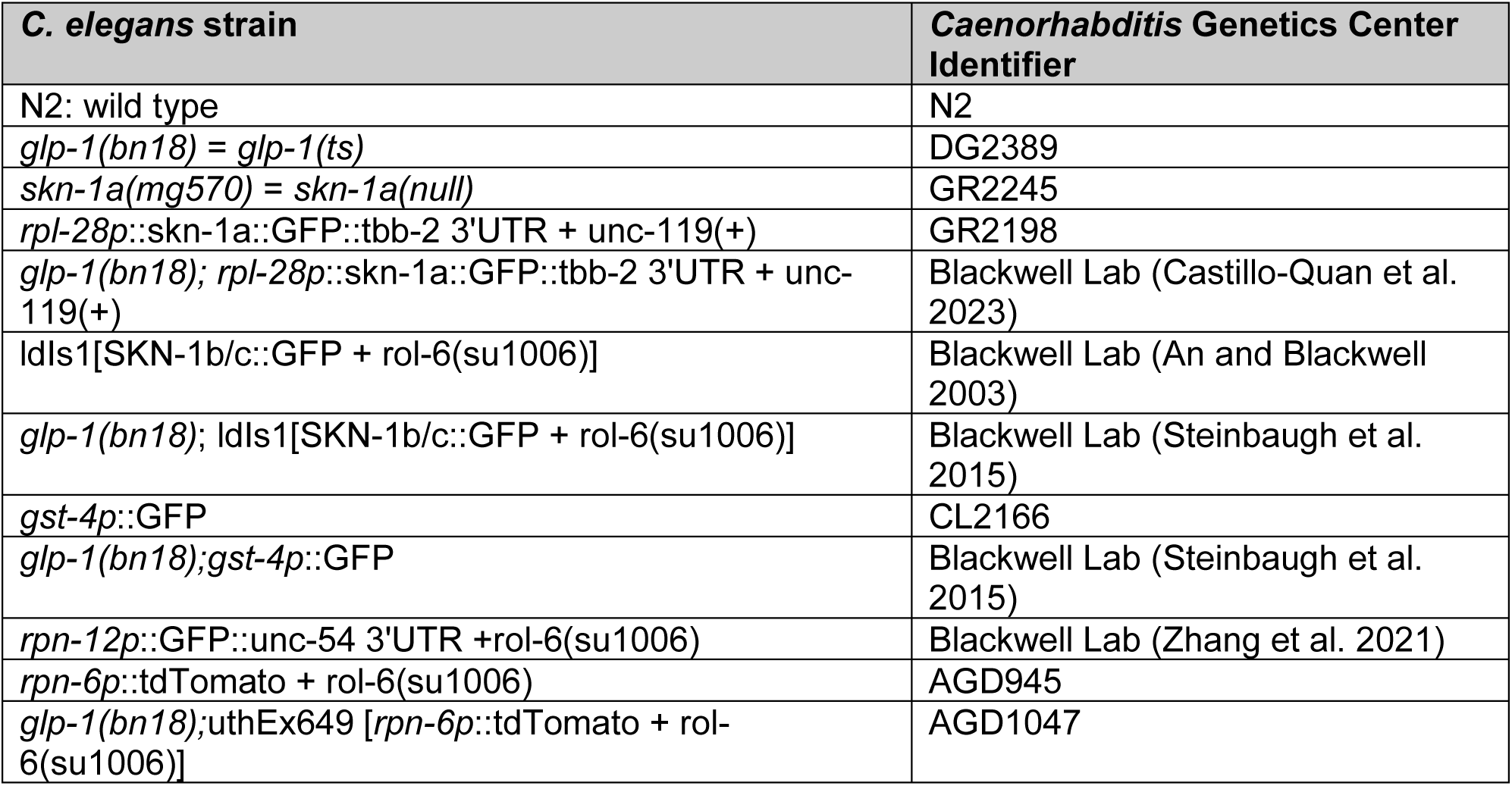

| *C. elegans* strain | *Caenorhabditis* Genetics Center Identifier |
| --- | --- |
| N2: wild type | N2 |
| *glp-1*(bn18) = *glp-1*(ts) | DG2389 |
| *skn-1a*(mg570) = *skn-1a*(null) | GR2245 |
| *rpl-28p*::*skn-1a*::GFP::tbb-2 3'UTR + unc-119(+) | GR2198 |
| *glp-1*(bn18); *rpl-28p*::*skn-1a*::GFP::tbb-2 3'UTR + unc-119(+) | Blackwell Lab (Castillo-Quan et al. 2023) |
| Idls1[SKN-1b/c::GFP + rol-6(su1006)] | Blackwell Lab (An and Blackwell 2003) |
| *glp-1*(bn18); Idls1[SKN-1b/c::GFP + rol-6(su1006)] | Blackwell Lab (Steinbaugh et al. 2015) |
| *gst-4p*::GFP | CL2166 |
| *glp-1*(bn18); *gst-4p*::GFP | Blackwell Lab (Steinbaugh et al. 2015) |
| *rpn-12p*::GFP::unc-54 3'UTR + rol-6(su1006) | Blackwell Lab (Zhang et al. 2021) |
| *rpn-6p*::tdTomato + rol-6(su1006) | AGD945 |
| *glp-1*(bn18); uthEx649 [*rpn-6p*::tdTomato + rol-6(su1006)] | AGD1047 |

### Bacterial and RNAi protocols

*C. elegans* were maintained on *E. coli* (OP50) bacteria as their standard food source. OP50 were cultured overnight in LB with streptomycin (10 mg/L) at 37°C. One milliliter of the liquid culture was seeded on standard NGM plates (10 cm petri dish) and air-dried at room temperature before use. RNAi was performed by feeding *C. elegans* HT115 bacteria, using the pL4440 plasmid as the empty vector (EV) control. The HT115 bacteria were grown using the standard RNAi techniques (Kamath and Ahringer 2003). Briefly, RNAi cultures were grown overnight at 37°C with shaking at 220 rpm in 50 mL conical tubes with 5 mL LB medium containing carbenicillin (50 mg/mL) and tetracycline (50 mg/mL). Cultures were then diluted 1:5 in LB containing carbenicillin and tetracycline for 6 hours to allow for reentry into the logarithmic growth phase. Cultures were then centrifuged at 3000 rpm for 10 min, and bacteria were resuspended in 5 ml of LB containing carbenicillin (50 mg/mL), tetracycline (50 mg/mL), and 1 mM isopropyl-β-D-thiogalactopyranoside (IPTG) (0.2 g/liter) before seeding onto NGM plates containing carbenicillin (50 mg/ml), tetracycline (50 mg/ml), and IPTG (0.2 g/liter). For double RNAi, clones were grown separately, in parallel, and after spin down, equal amounts (normalized by OD) of two clones or one clone and L4440 EV control were mixed and spread on plates. Bacterial clones that were used in experimental analysis were sequence-confirmed. RNAi was started at the L1 stage for all experiments except for longevity in which they were started at L4 stage.

#### Transcriptional GFP reporter analysis

Expression of *gst-4p*::GFP was used as a transcriptional reporter of SKN-1A and SKN-1C (Steinbaugh et al. 2015; Castillo-Quan et al. 2023), while *rpn-6p*::tdTomato and *rpn-12p*::GFP (Vilchez et al. 2012; Zhang et al. 2021; Castillo-Quan et al. 2023) were used as transcriptional reporters of the proteasome recovery response. Strains were maintained during development as described above. On day one of adulthood, animals were photographed and quantified. Scoring was as follows: low, no GFP or up to one-third of intestinal cells with GFP; medium, up to two-thirds of intestinal cells showed bright GFP or all intestinal cells showed dim GFP; high, two-thirds to all intestinal nuclei showed bright GFP. Chi-square test was used for statistical analyses using GraphPad Prism 9 (GraphPad Software, La Jolla, CA). For images, animals were immobilized for 5 min in 0.06% tetramisole/M9 buffer and aligned on nonseeded (empty) NGM plates. Images were taken with an Olympus IX51 microscope and cellSens standard 1.12 software.

#### Translational reporter intestinal nuclear localization

SKN-1A and SKN-1B/C translational reporter strains were grown, as described above, until day one of adulthood. At day one, animals were photographed and quantified. Animals were immobilized in 0.06% tetramisole/M9 buffer, mounted on 2% agarose pads on glass slides under coverslips, and imaged with ZEN 2012 software on an Axio Imager.M2 microscope (Zeiss, Jena, Germany). Scoring was as follows: low, no GFP or up to one-third nuclei contained GFP; medium, up to two-thirds of nuclei showed GFP; high, two-thirds to all intestinal nuclei showed GFP. In figures, the numbers above bars denote the sample size (biological replicates). Chi-square test was used for statistical analyses using GraphPad Prism 9 (GraphPad Software, La Jolla, CA).

#### Longevity Assays

Lifespan assays were completed using standard protocols (Castillo-Quan et al. 2023). Assays involving *glp-1(ts)* animals were performed at the nonpermissive temperature of 25°C, while all other lifespans were performed at 20°C. For all life-span experiments, *C. elegans* were L1 bleach-synchronized and seeded onto the appropriate bacteria. Animals were transferred at mid to late L4 stage to fresh plates containing 5′-fluorodeoxyuridine (FUdR) at a concentration of 50 µM to inhibit progeny development (Mitchell et al. 1979; Castillo-Quan et al. 2023). For RNAi experiments, bacteria was prepared as described above, and *C. elegans* were transferred at the L4 stage to RNAi plates containing FUdR. For *glp-1(ts)* experiments, the growth protocol and temperature shift were as described above. *C. elegans* were maintained at a density of 25 to 35 *C. elegans* per 6-cm plate on live bacteria and scored every other day. Animals that crawled off the plate, ruptured, or died from internal hatching were censored. Life spans were graphed as Kaplan-Meier survival curves, and analysis of survival curves was generated using log-rank test as previously described (Castillo-Quan et al. 2023).

#### Proteasomal stress and imaging fluorescence quantification

Proteasomal stress was induced with bortezomib (LC Laboratories, B-1408), a proteasome inhibitor as previously reported (Castillo-Quan et al. 2023). A 50mM stock of bortezomib was initially made, diluted in dimethyl sulfoxide. This stock was further diluted in M9 to 25μM. L4 stage animals were exposed to bortezomib by adding 600 μL of the 25μM bortezomib solution to the plate. 24 hours later fluorescence imaging and quantification was performed with an Olympus IX51 microscope and cellSens standard 1.12 software. Chi-square test was used for statistical analyses using GraphPad Prism 9 (GraphPad Software, La Jolla, CA).

#### Arsenite stress

For arsenite survival assay, day one adults were incubated in M9 buffer containing 5 mM AS (Riedel-de Haen, Seelze, Germany) and scored for survival hourly. Survival assays were graphed as Kaplan–Meier survival curves in Excel as previously described (Castillo-Quan et al. 2023). For arsenite transcriptional response using *gst-4p*::GFP synchronized L4 worms were washed off plates with M9 and incubated in 15 mL conical tubes containing 5 mM AS for four hours, rocking, at 25 °C. Afterwards animals were washed in M9 four times and pipette back onto fresh RNAi plates conditions for recovery. The following day, day-one adults were photographed and quantified. Chi-square test was used for statistical analyses using GraphPad Prism 9 (GraphPad Software, La Jolla, CA). For images, animals were immobilized for 5 min in 0.06% tetramisole/M9 buffer and aligned on non-seeded (empty) NGM plates. Images were taken with an Olympus IX51 microscope and cellSens standard 1.12 software.

#### Oil-Red-O TAG Staining

Oil-red O (ORO; Sigma-Aldrich, O0625-25G) staining was used to detect neutral lipids, of which *C. elegans* have only TAGs (O’Rourke et al. 2009; Watts and Ristow 2017; Castillo-Quan et al. 2023). ORO staining was performed on fixed animals as previously described (Steinbaugh et al. 2015; Castillo-Quan et al. 2023). Five hundred to one thousand L1 bleach-synchronized *C. elegans* were grown to adulthood day one and washed three times with phosphate-buffered saline (PBS). Upon the last wash, animals were treated with PBS containing 2% paraformaldehyde (PFA; Santa Cruz Biotechnology, SC-281692) and then snap-frozen at least four times in a dry ice/ethanol bath to permeabilize the cuticle. *C. elegans* samples were then washed at least twice with PBS. They were then washed twice with a 60% isopropanol solution to remove the PFA. After removal of the isopropanol solution, samples were treated with freshly prepared and filtered ORO solution (0.5 g of ORO powder in 100 ml of 60% isopropanol). Samples were then stained overnight in 1.5 mL Eppendorf tubes at room temperature with gentle shaking. Animals were imaged at (40x magnification) using differential interference contrast microscopy. The region anterior of the worm’s embryos was centered in images acquired. Images were adjusted to a standard Red, Green, Blue value (RGB) to maximize red appearance. RGB settings were retained identically throughout individual replicates. TIF images were entered into ImageJ and the split channels function was used to isolate the greatest red signal; the same file type (either Red, Green, or Blue) was used in all images in individual replicates. The subtract background function was then used on all images. The lower manual threshold was set to zero, and the upper manual threshold was set to five. The freehand selection function was used to select the upper mid intestine region, and then the measure function was used. Values were then standardized to *C. elegans* in empty vector (EV) RNAi using Excel. Statistical analysis was performed by ANOVA with Tukey post hoc analysis using GraphPad Prism 9.

#### Quantitative reverse transcription polymerase chain reaction

Samples were prepared from 1500 day-1 adult bleach-synchronized *C. elegans*, as previously described (Castillo-Quan et al. 2023). Briefly, RNA was extracted using TRIzol-based (Thermo Fisher Scientific, 15596026) phenol-chloroform extraction and purified with RNA Clean and Concentrator-5 spin columns (Zymo Research, R2050). RNA concentration and quality were assessed with a NanoDrop 1000 spectrophotometer (Thermo Fisher Scientific). Complementary DNAs were prepared using the SuperScript III First-Strand Synthesis SuperMix for qRT-PCR (Thermo Fisher Scientific). mRNA levels were quantified from biological triplicates and technical duplicates-triplicates using SYBR Green (Thermo Fisher Scientific, 11760500) fluorescence on a 384-well format real-time PCR 7900 (Applied Biosystems, Foster City, CA). After an initial denaturation step (95°C for 10 min), amplification was performed using 40 cycles of denaturation (95°C for 15 s) and annealing (60°C for 1 min). Samples were analyzed by the standard curve method, with normalization two reference genes (Castillo-Quan et al. 2023). *P* values were calculated by ANOVA with Tukey post hoc analysis in GraphPad Prism 9. The primers used in this study are listed here:

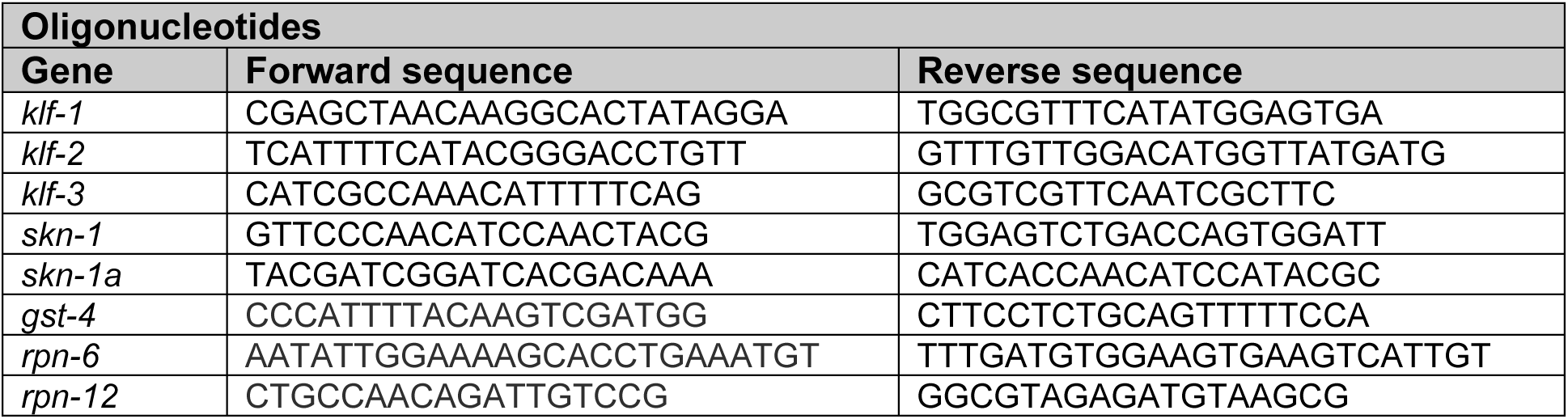
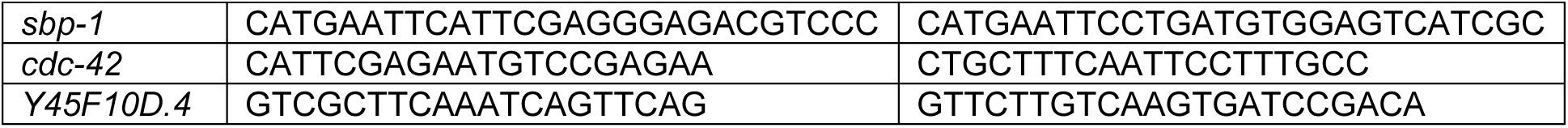

## Acknowledgments

We would like to dedicate this work to the loving memory of our dearest mentor and esteemed colleague Dr. T. Keith Blackwell (1957-2025), who was instrumental in the development of this project. Dr. Blackwell is widely recognized as a world leader in SKN-1 biology. We thank Blackwell lab members and Emmanuel College Biology faculty for helpful discussions.

## Author contributions

Conceptualization: JICQ, TKB, NM.

Investigation: JICQ, AM, UK, KG, AC, JM, JB, MLT, EJ, AC, NM.

Supervision: JICQ, TKB, JES, NM.

Writing original draft: JICQ, NM.

Writing, review & editing: JICQ, JES, NM.

---

**Figure S1.**
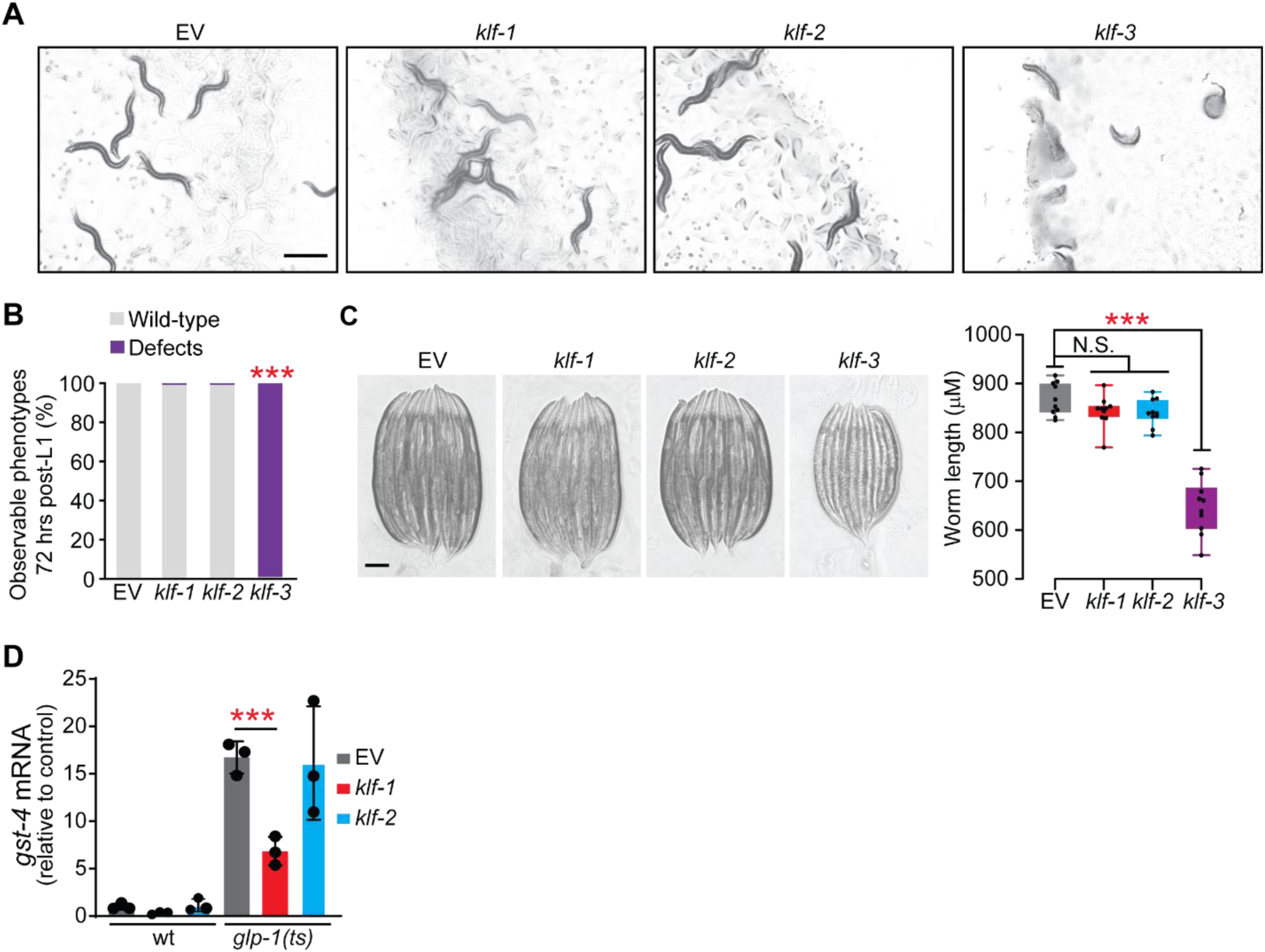
KLFs role in development and SKN-1 activity. **(A)** Images of adult day one animals after exposure to different RNAi from L1. Images were taken on growth plates 72 hours post L1. Scale bar, 500 μm. Knockdown of *klf-3* results in gross developmental defects, observed through a dissecting stereoscope. **(B)** Quantification of developmental defective phenotypes in a plate of 100 animals. Phenotypes quantified: uterine prolapse, intestinal prolapse, bagging, and smaller size. Pairwise Chi-square tests with False Discovery Rate (FDR) correction (via Benjamini-Hochberg) for multiple comparisons. **(C)** Size was quantified by measuring worm length using Image J. Representative images (left) and quantification (right). ANOVA with Tukey multiple comparisons test. **(D)** GSC(−) animals have higher expression of *gst-4* mRNA compared to wild type animals, with *klf-1* knockdown significantly reducing it. ANOVA with Tukey multiple comparisons test. Error bars represent standard deviation.

**Figure S2.**
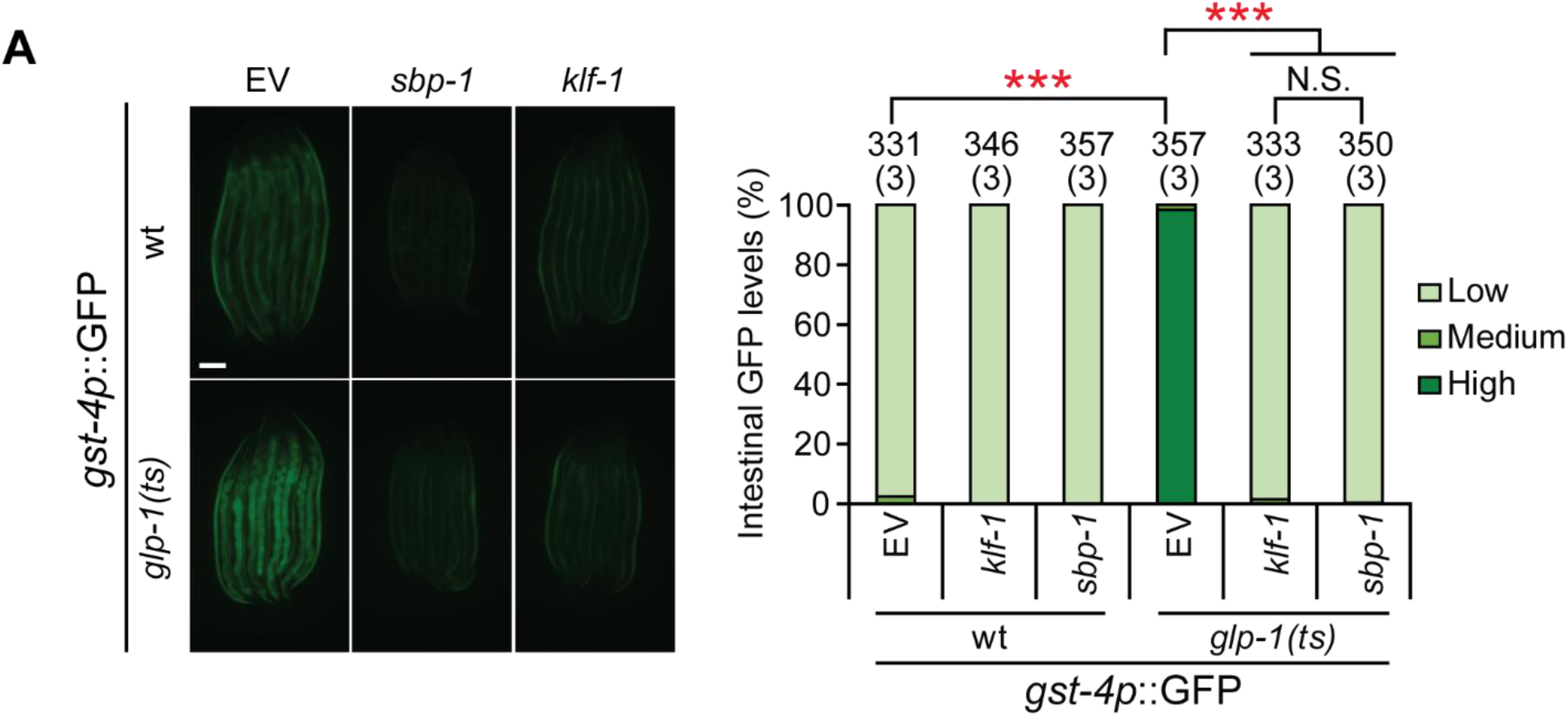
The role of KLF-1 and SBP-1 on SKN-1 transcription. **(A)** Increased expression of *gst-4p*::GFP in GSC(−) animals is similarly reduced by knockdown of *klf-1* or *sbp-1*/SREBP1. Representative images (left) and quantification (right; see data S2 for details). Scale bar, 200 μm. Numbers above bars denote sample size (biological replicates). Pairwise Chi-square tests with False Discovery Rate (FDR) correction (via Benjamini-Hochberg) for multiple comparisons. N.S. = not significant; \**P* < 0.05, ***p < 0.0001.

**Figure S3.**
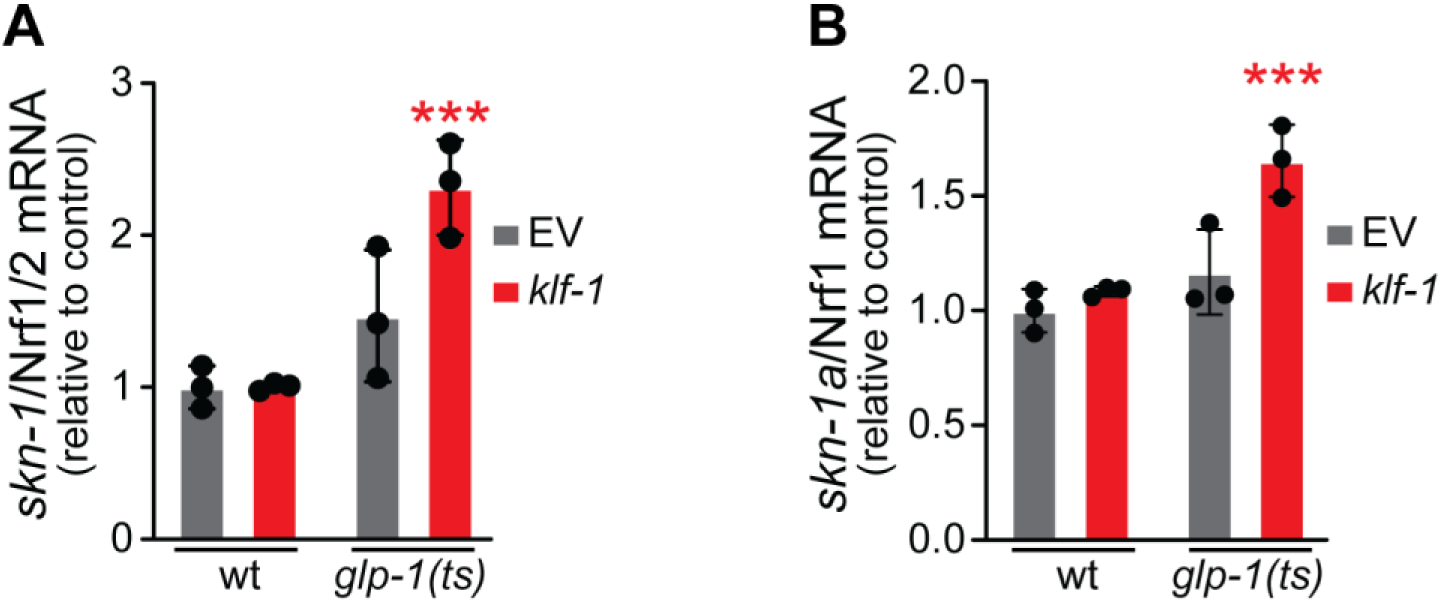
KLF-1 does not affect *skn-1* transcription. **(A)** mRNA levels of *skn-1* (all isoforms) are unaffected by GSC(−), and significantly higher in GSC(−) animals after they undergo *klf-1* knockdown. **(B)** Similarly, mRNA levels of specifically *skn-1a* are unchanged by GSC(−), but increase in GSC(−) animals subjected to *klf-1* RNAi. Representative images (left) and quantification (right; see data S2 for details). Scale bar, 200 μm. ANOVA with Tukey multiple comparisons test. Error bars represent standard deviation. ***p < 0.0001.

**Figure S4.**
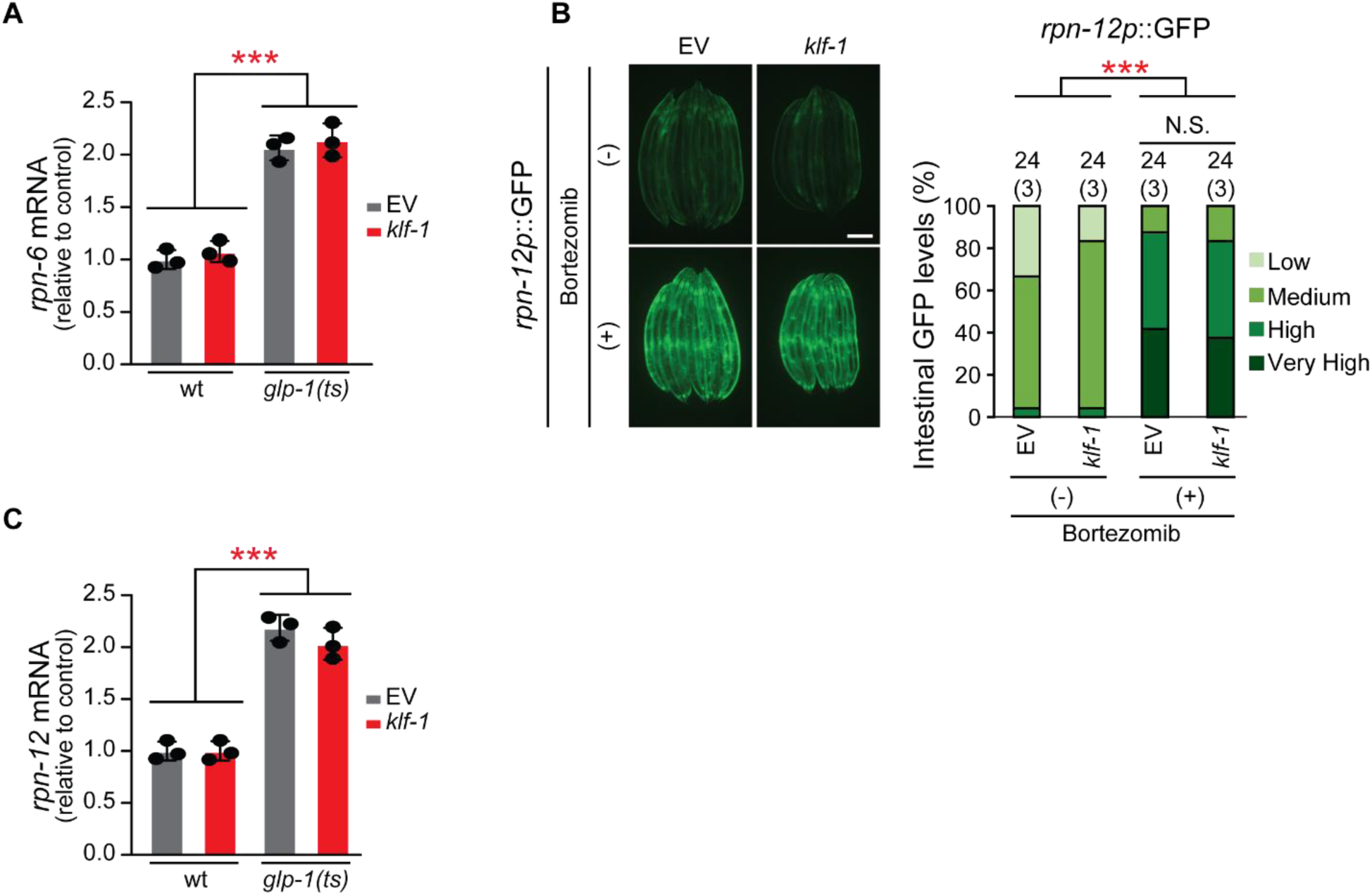
KLF-1 does not modify proteasome subunit gene expression or the proteasome recovery response. **(A)** *rpn-6* mRNA levels increase in GSC(−) animals and do not change upon *klf-1* knockdown. ANOVA with Tukey multiple comparisons test. Error bars represent standard deviation. ***p < 0.0001. **(B)** Bortezomib increases *rpn-12p*::GFP expression in the canonical SKN-1A proteasome recovery response, while *klf-1* knockdown does not have an effect. Numbers above bars denote sample size (biological replicates). Pairwise Chi-square tests with False Discovery Rate (FDR) correction (via Benjamini-Hochberg) for multiple comparisons. **(C)** *rpn-12* mRNA levels increase in GSC(−) animals and do not change upon *klf-1* RNAi. ANOVA with Tukey multiple comparisons test. Error bars represent standard deviation. ***p < 0.0001.

**Figure S5.**
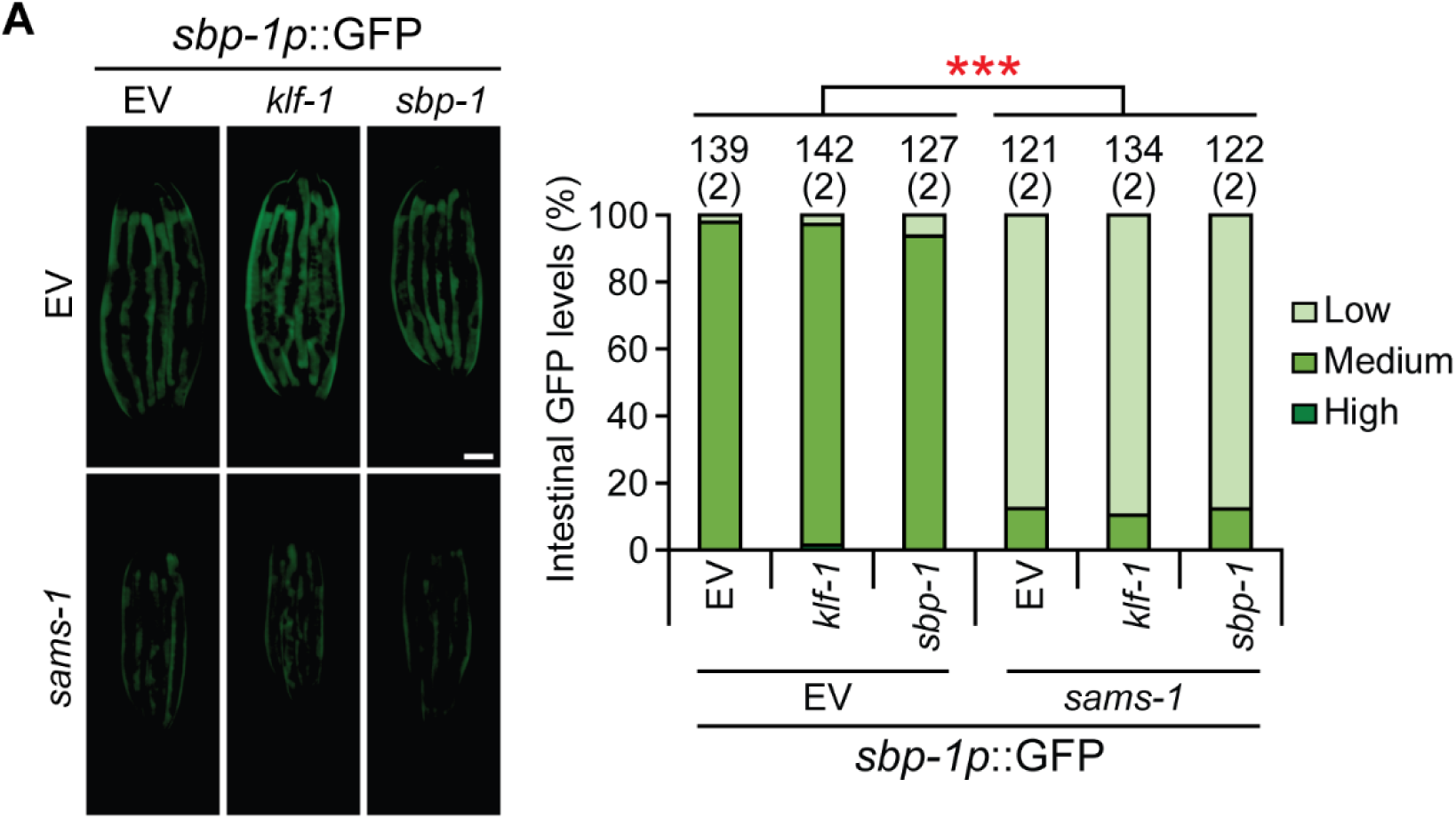
KLF-1 does not affect *sbp-1* transcription. **(A)** *sbp-1p*::GFP transcriptional reporter expression is not significantly affected by *klf-1* knockdown. *sams-1* knockdown, which activates SBP-1, mildly but significantly reduces *sbp-1* transcriptional reporter expression, potentially as a compensatory response. Representative images (left) and quantification (right; see data S2 for details). Scale bar, 200 μm. Numbers above bars denote sample size (biological replicates). False Discovery Rate (FDR) correction (via Benjamini-Hochberg) for multiple comparisons. ***p < 0.0001.

